# BoneMA – Synthesis and Characterization of a Methacrylated Bone-derived Hydrogel for Bioprinting of Vascularized Tissues

**DOI:** 10.1101/2020.03.02.974063

**Authors:** S. Prakash Parthiban, Avathamsa Athirasala, Anthony Tahayeri, Reyan Abdelmoniem, Anne George, Luiz E. Bertassoni

**Author notes:** These authors contributed equally to this work (S. Prakash Parthiban, Avathamsa Athirasala).

## Abstract

It has long been proposed that recapitulating the extracellular matrix (ECM) of native human tissues in the laboratory may enhance the regenerative capacity of engineered scaffolds in-vivo. Organ- and tissue-derived decellularized ECM biomaterials have been widely used for tissue repair, especially due to their intrinsic biochemical cues that can facilitate repair and regeneration. The main purpose of this study was to synthesize a new photocrosslinkable human bone-derived ECM hydrogel for bioprinting of vascularized scaffolds. To that end, we demineralized and decellularized human bone fragments to obtain a bone matrix, which was further processed and functionalized with methacrylate groups to form a photocrosslinkable methacrylate bone ECM hydrogel – BoneMA. The mechanical properties of BoneMA were tunable, with the elastic modulus increasing as a function of photocrosslinking time, while still retaining the nanoscale features of the polymer networks. The intrinsic cell-compatibility of the bone matrix ensured the synthesis of a highly cytocompatible hydrogel. The bioprinted BoneMA scaffolds supported vascularization of endothelial cells and within a day led to the formation of interconnected vascular networks. We propose that such a quick vascular network formation was due to the host of pro-angiogenic biomolecules present in the bone ECM matrix. Further, we also demonstrate the bioprintability of BoneMA in microdimensions as injectable ECM-based building blocks for microscale tissue engineering in a minimally invasive manner. We conclude that BoneMA may be a useful hydrogel system for tissue engineering and regenerative medicine.

## 1. Introduction

The rational design of biomaterials for regenerative applications has long sought to approximate the biological composition of the native extracellular matrix (ECM). The main components of the human ECM are collagens, proteoglycans, laminin, fibronectin, and elastin, which, along with other matrix macromolecules and growth factors, are linked together to form a structurally stable network that contributes to the mechanical properties of different tissues [1]. The ECM, however, is tissue-specific, where cells secrete matrix molecules based on their local conditions, such as biological function, mechanical loading, hypoxia, and variability in nutrient concentration [2–4]. Furthermore, the composition of the ECM varies dynamically through life to regulate various processes of development, differentiation, and remodeling [5–7]. The bioactive nature of the native ECM of various tissues has been well established [7, 8]. Therefore, researchers have capitalized on the cell instructive and regenerative potential of these naturally bioactive materials to develop matrix-derived scaffolds for regenerative applications, examples of which include repair of bone [9, 10], cartilage [11], brain and spinal cord [12, 13], pancreas [14] and a host of other tissues [15].

Despite the remarkable capacity of ECM-derived proteins to facilitate tissue regeneration and healing, and their inherent capacity to self-assemble into structured hydrogels, the primary disadvantage of these unmodified native materials is the lack of control over their material properties, such as stiffness, degradation, porosity, etc. All of these properties have been demonstrated to significantly influence cell behavior. Additionally, their weak polymerization chemistry and resultant material properties do not support the fabrication of 3D constructs with the complex microarchitectural features that are typically associated with native tissues. In order to address these limitations in the context of bone-derived hydrogels, here we developed a novel methacrylated bone-derived biomaterial (BoneMA) consisting of bone ECM proteins functionalized with methacrylates, which render the material photocrosslinkable in the presence of a photoinitiator, while maintaining the biological advantages associated with the composition of the native ECM. We hypothesized that similar to other popular photocrosslinkable biomaterials, such as gelatin methacryloyl (GelMA) [16, 17] or methacrylated hyaluronic acid (MeHA) [18], photopolymerization of bone ECM proteins using these light sensitive moieties would enable formation of hydrogel scaffolds with precisely tuned and reproducible microscale architectures and physico-mechanical properties, through modulation of light exposure. Furthermore, the photocrosslinkable nature of this material should allow for straightforward bioprinting of cell-laden micro-tissue constructs using a digital light processing (DLP) based approach, which is advantageous over conventional methods of fabrication of demineralized and de-cellularized bone [19].

To illustrate the applicability of BoneMA hydrogels as a biomaterial for tissue engineering and bioprinting, here we demonstrate the cytocompatibility of this novel material system through cell viability assays, following which, we assessed vascular network formation by encapsulated human umbilical vein endothelial cells (HUVEC) after printing. We also compare the material’s vasculogenic capacity in comparison with GelMA – a methacrylated biomaterial of known vasculogenic potential – and finally, we examined the potential application of these scaffolds as an injectable pre-vascularized micro tissue constructs with a range of bioprinted microgeometries.

## 2. Experimental

### 2.1. Preparation of demineralized and decellularized bone matrix

Demineralized and decellularized ECM bone matrix was extracted according to previously published protocols [19]. In short, fresh human osteochondral/osteoarticular bone fragments were scraped thoroughly to remove residual tissue and rinsed with Dulbecco’s phosphate buffered saline (DPBS) solution containing penicillin (100 IU/ml) and streptomycin (100 μg/ml) (Corning, USA). The bone fragments were sectioned using a low-speed diamond saw (Struers Accutom-5) and the resulting sections were then ground to a fine powder using a cryogenic freezer/mill (SPEX 6700, USA). The finely ground bone powder was demineralized under agitation using 0.5 N HCl (25 ml per gram) for 24 hours and the insoluble portion of the bone matrix was retrieved by vacuum-filtration after rinsing with distilled water. After lipid removal by solvent extraction with 1:1 (v/v) chloroform (Sigma-Aldrich, USA)-methanol (Acros Organics, USA) solution for 1 hour, followed by rinsing with methanol and distilled water, the demineralized bone matrix was flash-frozen in liquid nitrogen and lyophilized overnight (Labconco, USA). Subsequently, the demineralized bone matrix was decellularized with 0.05% trypsin-0.02% ethylenediaminetetraacetic acid (EDTA) at 37 °C, 5% CO2 under continuous agitation for 24 hours and then rinsed with DPBS containing penicillin (100 IU/ml) and streptomycin (100 μg/ml). Following this, the processed bone matrix was subjected to enzymatic digestion by pepsin (1 mg/ml in 0.01 N HCl), where a suspension of 10 mg of matrix per ml of pepsin was stirred under agitation for 96 hours until the matrix was solubilized. Finally, the solubilized matrix was centrifuged for 30 min at 4 °C, filtered under vacuum and stored at −20 °C.

### 2.2. Synthesis of BoneMA

For the synthesis of BoneMA, the frozen demineralized and decellularized bone matrix was lyophilized for two days. Next, 10% (w/v) of the lyophilized matrix was dissolved in 0.25 M carbonate-bicarbonate buffer, and the pH was adjusted to 9 using 4 M NaOH or 6 M HCl under constant stirring at 50°C. Once the bone matrix was dissolved, methacrylic anhydride (0.2 ml/g of bone matrix) was added to the solution in a dropwise manner under constant stirring while maintaining the temperature. The reaction was allowed to proceed for 3 hours, and the pH was maintained at 9 by adding 4 M NaOH. After the completion of the reaction, the modified bone matrix solution was diluted four times with warm deionized water. Unreacted methacrylic anhydride was removed from the solution by dialyzing it against deionized water for 24 hours, following which the solution was filtered, lyophilized and stored at −80 °C until further use (Figure 1A). In order to confirm the methacrylation of the bone matrix, we performed nuclear magnetic resonance (NMR) on the demineralized bone matrix and BoneMA using a Bruker 400 MHz Avance II+ spectrometer with a magnetic field strength of 400 MHz and chloroformd as the solvent. The NMR spectra readout of the reaction product was compared with that of unmodified bone ECM as a control.

**Figure 1.**
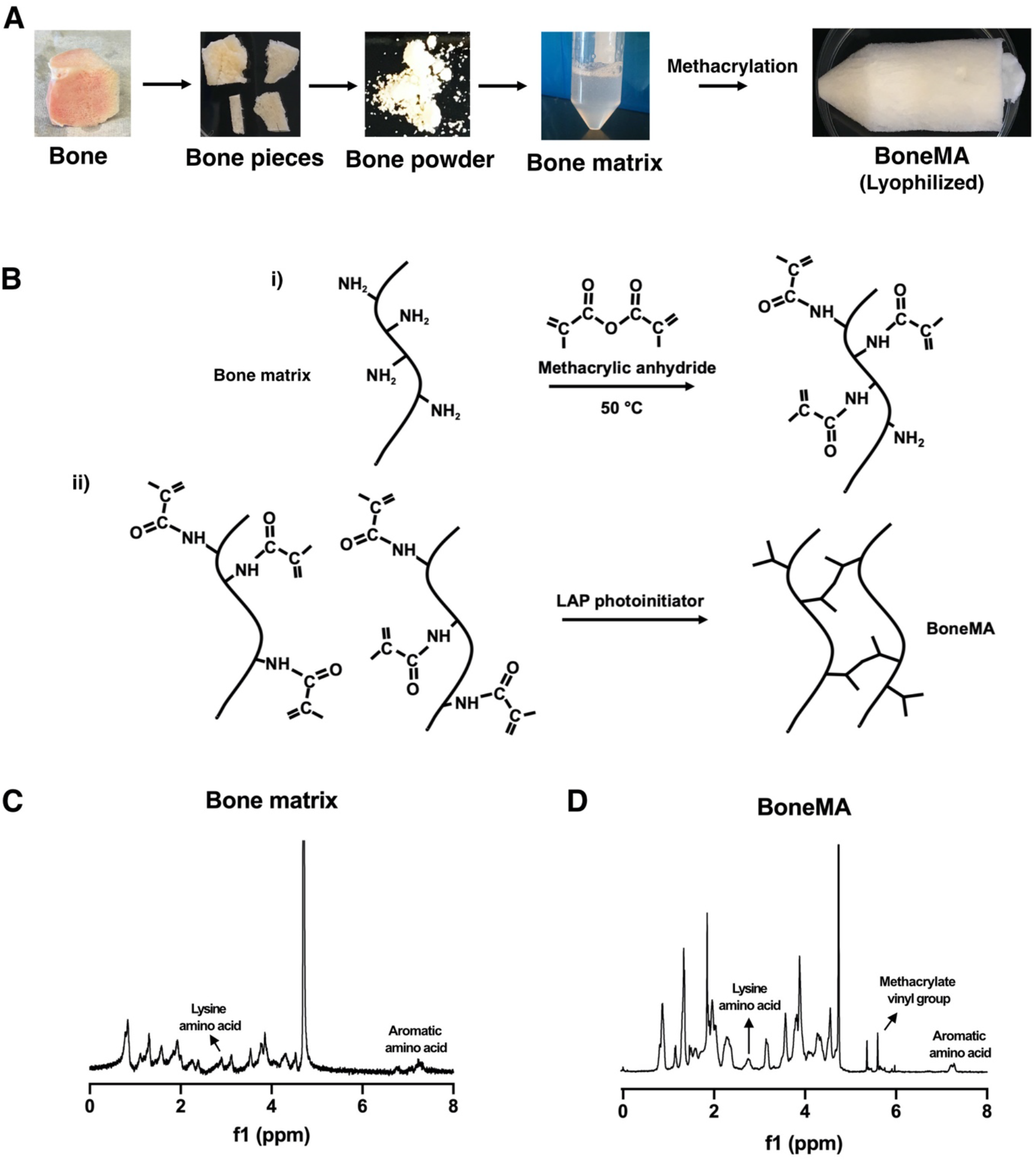
Synthesis and chemistry of BoneMA. (A) Schematic depicting the synthesis of BoneMA wherein osteochondral/osteoarticular bone were powdered, demineralized, and processed for removal of lipids and decellularization procedures to extract bone matrix proteins. The processed bone matrix was then modified by methacrylation to form BoneMA macromers (B) through i) an addition reaction with methacrylic anhydride where methacrylate groups are linked to the pendant amine groups. ii) BoneMA is polymerized by visible light in the presence of LAP photoinitiator through crosslinking of these methacrylate groups to form BoneMA hydrogel. Comparison of the NMR spectra of (C) unmodified bone matrix and (D) BoneMA showed a peak occurring between 5.3 and 6.3 ppm representing the vinyl group of methacrylates in BoneMA, in addition to the proton signal for aromatic amino acid and lysine groups in the bone matrix, thereby confirming derivatization of bone matrix to form BoneMA.

### 2.3. Sample preparation

The freeze-dried BoneMA was dissolved in DPBS (5% (w/v)) with 0.15 w/v% Lithium phenyl-2,4,6-trimethylbenzoylphosphinate (LAP, Tokyo Chemical Industry, L0290) photoinitiator. A digital light processing (DLP) 3D printer (Ember, Autodesk) was used to prepare hydrogel samples for different experiments by exposing the BoneMA hydrogel prepolymer to 20 mW/cm^2^ of light (405 ±5 nm) for 15, 30, and 45 sec. For physical and mechanical characterization, samples were printed with 5 mm in diameter and 1.5 mm in thickness, whereas biological characterization tests used 450 thick specimens printed with geometrical shapes described below (N=5).

### 2.4. Mechanical properties

The hydrogel mechanical properties were measured using a DHR-1 rheometer (TA Instruments) fitted with a UV curing accessory containing a bandpass filter for 405 ±5 nm. The intensity of the light from an Acticure® 4000 (EXFO) coupled with the rheometer was matched to the DLP printer at 20 mW/cm^2^ using a power meter (Power Max5200, Molectron). Shear modulus measurements were performed as a function of crosslinking time under 0.1% strain and frequency of 10 Hz and were converted to elastic modulus using a previously reported equation [20].

### 2.5. Scanning electron microscopy (SEM)

The BoneMA hydrogel samples were prepared according to the above procedure for all time points. For SEM imaging, the BoneMA hydrogel was immersed in 5 mL DPBS at 37 °C for 24 h (n = 5). Next, samples were fixed with 2.5% glutaraldehyde solution for 1 hour. After fixing, the samples were dehydrated for 10 minutes in a series of ethanol solutions of concentration 25%, 50%, 75%, 90%, and 100%, respectively. After dehydration, samples were critical point dried for 3 hours. These dried samples were coated with gold/palladium and imaged with FEI Quanta 200 SEM at 15 kV.

### 2.6. Cell culture

Since different aspects and applications of BoneMA were being tested, samples were prepared using different cell lineages for various experiments. To that end, human dental pulp stem cells (HDPSC) and human mesenchymal stem cells (HMSC) were used to assess cytocompatibility and bioprintability, while green fluorescent protein (GFP)-expressing human umbilical vein endothelial cells (HUVECs) (cAP0001GFP, Angio-proteomie, USA) were employed to investigate the vasculogenic potential of the synthesized biomaterial. HDPSCs (Lonza, USA) and HMSCs were each cultured in a-MEM medium supplemented with 10% (v/v) fetal bovine serum and 1% (v/v) penicillin-streptomycin. HUVECs were cultured in Endothelial Cell culture medium (Vasculife-VEGF, Lifeline Cell Technologies) on 0.1% gelatin-coated substrates. Cells were maintained in an incubator at 37°C, 5% CO2 incubator, and the medium was replaced every 2 days.

### 2.7. Cell viability

In order to assess the cytocompatibility of BoneMA, HDPSCs were trypsinized and resuspended in a BoneMA precursor (5% (w/v)) containing LAP (0.15% (w/v)) at a cell density of 5 x 10^5^ cells/mL. This cell-laden prepolymer solution was photopolymerized as described above for 15, 30, and 45 sec. Cell viability was measured on days 1, 3, and 7 post-encapsulation using a membrane permeability based fluorescent live/dead staining kit (Molecular Probes), and the fraction of live cells was estimated using ImageJ software from at least 3 distinct regions per sample (N=5).

### 2.8. Fabrication of vascularized hydrogels

The vasculogenic potential of BoneMA was examined in comparison to GelMA, which was synthesized with a comparable degree of methacrylation (supplementary methods S1) and photo-crosslinked for the same amounts of time. To that end, HUVECs were encapsulated in BoneMA hydrogels precursors (5% (w/v)) containing LAP (0.15% (w/v)) at a cell density of 1 x 10^7^ cells/ml, crosslinked for 15, 30 and 45 s as described previously. Vascular capillary network formation in these hydrogels was characterized daily for 7 days and quantified for the number of segments, number of end points and junctions, as well as the total length of the network using an ImageJ, as described previously [21].

### 2.9. Bioprinting of BoneMA geometries

To bioprint BoneMA in various geometrical shapes, specific print patterns were designed using CAD (Fusion 360, AutoDesk) and converted into image slices with the accompanying 3D printing software (Print Studio, Autodesk). The print patterns were designed arbitrarily with star, square, triangle, and rhombus shapes, as well as flower, spiral, concentric circles, and the OHSU logo. Shapes were printed with length and width dimensions that ranged from 600 μm (square) to 2.5 mm (OHSU logo) with a thickness of 450 μm, under printing exposures of 25 sec. For bioprinting of the said geometries, the bioink was formulated by mixing HMSCs with the BoneMA (5% (w/v)) pre-polymer containing LAP (0.15% (w/v)) at a concentration of 5 x 10^5^ cells/ml. The HMSC-laden bioprinted BoneMA was fixed with 4% (v/v) paraformaldehyde and permeabilized using 0.1% (v/v) Triton X-100. Next, the samples were treated with 1.5% (w/v) bovine serum albumin (Sigma Aldrich) in DPBS for 1h, followed by Image-iT FX signal enhancer (Invitrogen, CA) for 30 min, after which they were immunostained with ActinGreen™ 488 ReadyProbes™ (Invitrogen).

### 2.10. Statistical analysis

All data are presented as mean ±standard deviation. Data were compared using one-way ANOVA followed by Tukey posthoc test (a = 0.05) with Graphpad Prism 8.

## 3. Results

In order to verify the addition of methacrylates to the molecular structure of the bone matrix, we performed nuclear magnetic resonance analysis before and after the reaction with methacrylic anhydride. The spectra of unmodified bone matrix showed the expected peaks corresponding to the amino acids containing primary amines and aromatic groups (Figure 1C), whereas the BoneMA macromer had peaks between 5 and 6 ppm, indicative of the presence of the vinyl component of methacrylates (Figure 1D), thus confirming functionalization of the bone matrix proteins.

Next, the tunability of the mechanical properties of the newly formed methacrylated material was examined as a function of light exposure by comparing the elastic moduli of hydrogels polymerized for 15, 30 and 45 seconds, using the DLP 3D printer as a light source. The elastic modulus of BoneMA was found to increase significantly as a function of the duration of light exposure, starting at 0.9 kPa when crosslinked for 15 s, and increasing gradually to 1.3 (p<0.05) and 1.5 (p<0.001), at 30 and 45 s respectively (Figure 2A).

**Figure 2.**
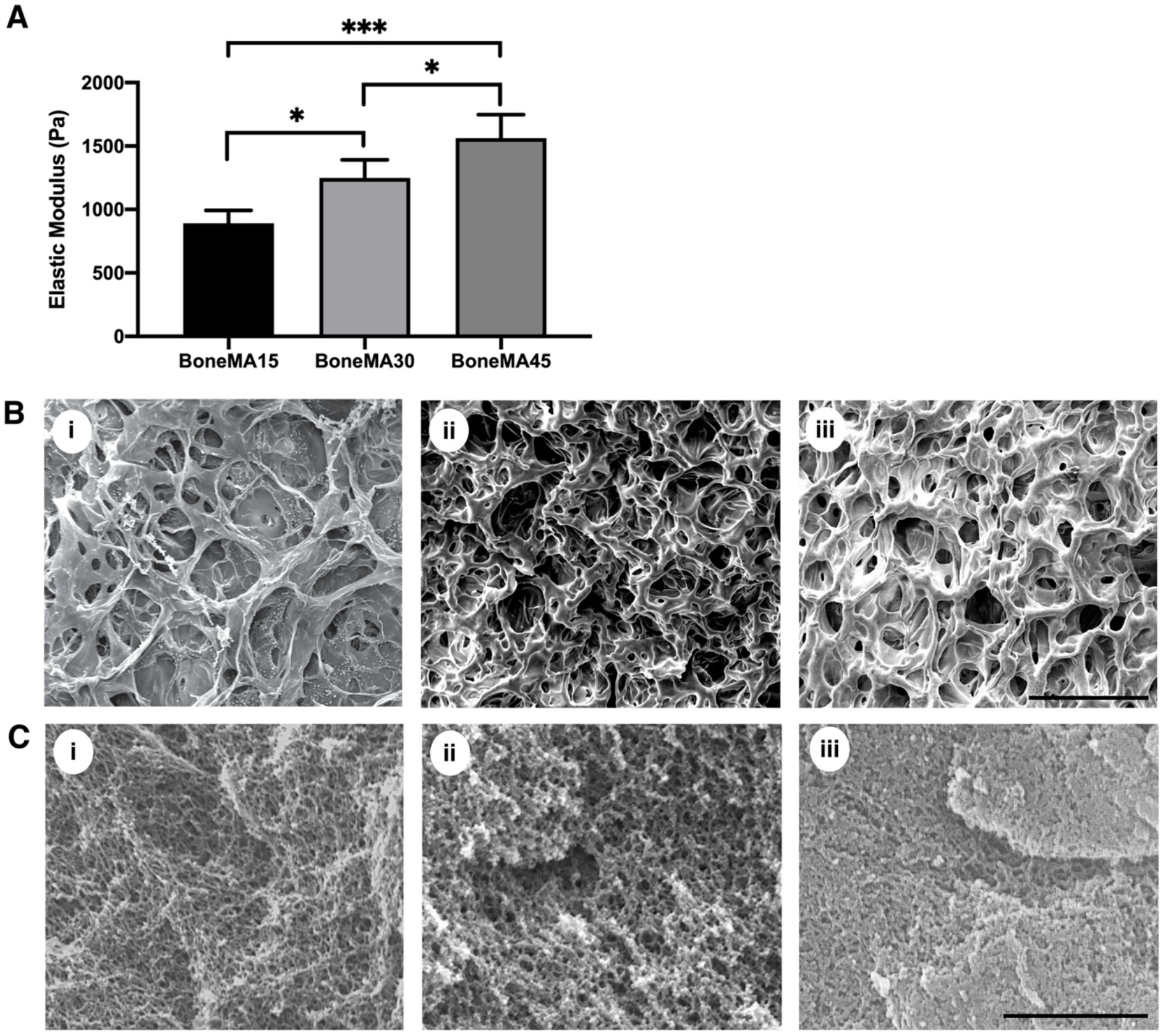
Physical characterization of BoneMA. The mechanical properties of the BoneMA were tunable by light exposure with (A) the elastic modulus increasing gradually as a function of crosslinking time from 15 to 30 and 45 sec (\**p* < 0.05; \*\*\**p* < 0.001). Similarly, the apparent pore size, as visualized by SEM images of (B) freeze-dried (Scale – 100 μm) and (C) critical point dried BoneMA samples (Scale – 2 μm), decreased with crosslinking duration from (i) 15 sec to (ii) 30 and (iii) 45 sec.

SEM images showed the presence of the typical pore-like microstructures that are commonly observed in covalently crosslinked hydrogels with BoneMA crosslinked for 15 seconds having distinctly larger pores than its more crosslinked counterparts (Figure 2B). Critical point drying of the samples enabled imaging of the highly interconnected fibrous micromorphology of the hydrogels (Figure 2C), suggesting that, despite the methacrylation of the solubilized and pepsinized fibrillar proteins in the bone matrix, a semi-fibrillar hydrogel still formed on the nanoscale. Notably, the same porous and fibrous morphology was observed in GelMA hydrogels (Supplementary Figure S1B, C). However, while there was no significant difference in the swelling mass ratio of BoneMA as a function of crosslinking time, dye permeability studies showed that the rate of diffusion through each of these BoneMA hydrogels mirrored their apparent pore size (Supplementary Figure S2).

HDPSCs remained highly viable through at least 7 days in culture within all bioprinted BoneMA hydrogel constructs, with average viability of 91, 94 and 90% for BoneMA crosslinked for 15, 30 and 45 seconds on day 7, respectively (Figure 3). These results confirmed the cytocompatibility of the biomaterial in comparison with more established hydrogel scaffolds, such as GelMA (Supplementary Figure S3), which was used as a control.

**Figure 3.**
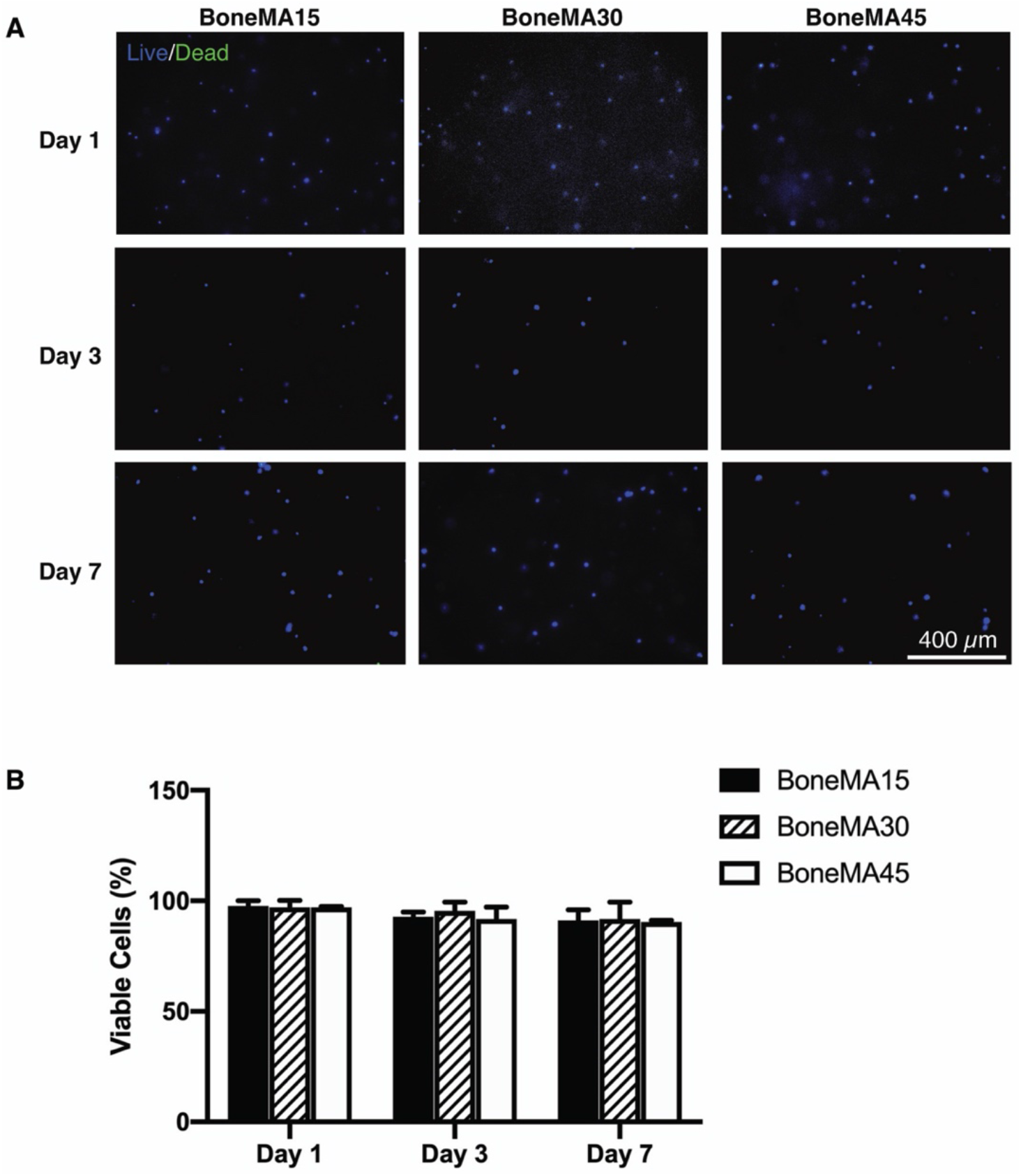
Cytocompatibility of BoneMA. (A) Representative images of HDPSCs stained for live (blue) and dead (green) cells on days 1, 3, and 7 after encapsulation in BoneMA hydrogels crosslinked for 15, 30 and 45 sec showed (B) very high cell viability across all groups and time points.

Next, in order to assess the vasculogenic potential of BoneMA, we measured vascular network formation in HUVEC-laden hydrogels crosslinked for 15, 30, and 45 sec (Figure 4A), and then compared it against similarly crosslinked GelMA hydrogels (Figure 4B), which was used as a control. Better vascular network formation was observed in the less crosslinked hydrogels at each time point in both BoneMA and GelMA suggesting that softer mechanics and higher porosity supported the formation of capillaries. However, while the network formation began in as little as 24 h in BoneMA hydrogels, with almost fully established capillary formation by day 2 in BoneMA hydrogels, network formation was much slower in GelMA, which only started around day 2 and was more established during day 3. Since chemoattraction for vascular network formation is innately dependent on the physico-mechanical properties of the hydrogel, especially the stiffness [22], we directly compared capillary formation in BoneMA and GelMA hydrogels of comparable elastic moduli. Interestingly, BoneMA crosslinked for 45 seconds to an average modulus of 1.56 kPa, showed faster network formation than and GelMA crosslinked for 15 seconds to a comparable modulus of 1.57 kPA (Figure 5).

**Figure 4.**
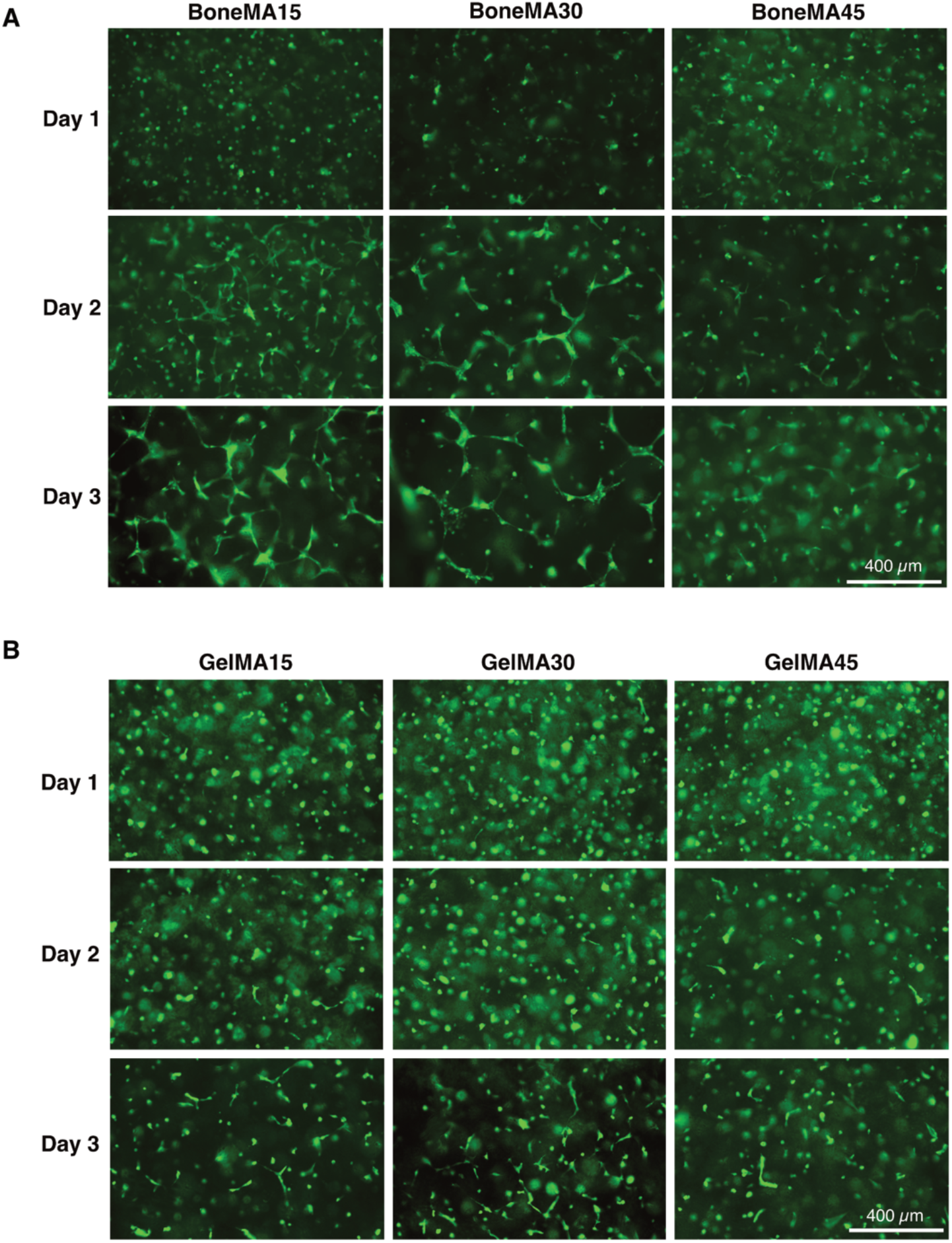
Vascular network formation in BoneMA and GelMA. Representative fluorescence images of GFP-HUVECs in (A) BoneMA and (B) GelMA hydrogels, each crosslinked for 15, 30, and 45 sec showed better vascular network formation in the less crosslinked hydrogels. Importantly, capillary formation started earlier and grew faster in BoneMA hydrogels in comparison to that in GelMA hydrogels crosslinked similarly. (Scale bar – 400 μm).

**Figure 5.**
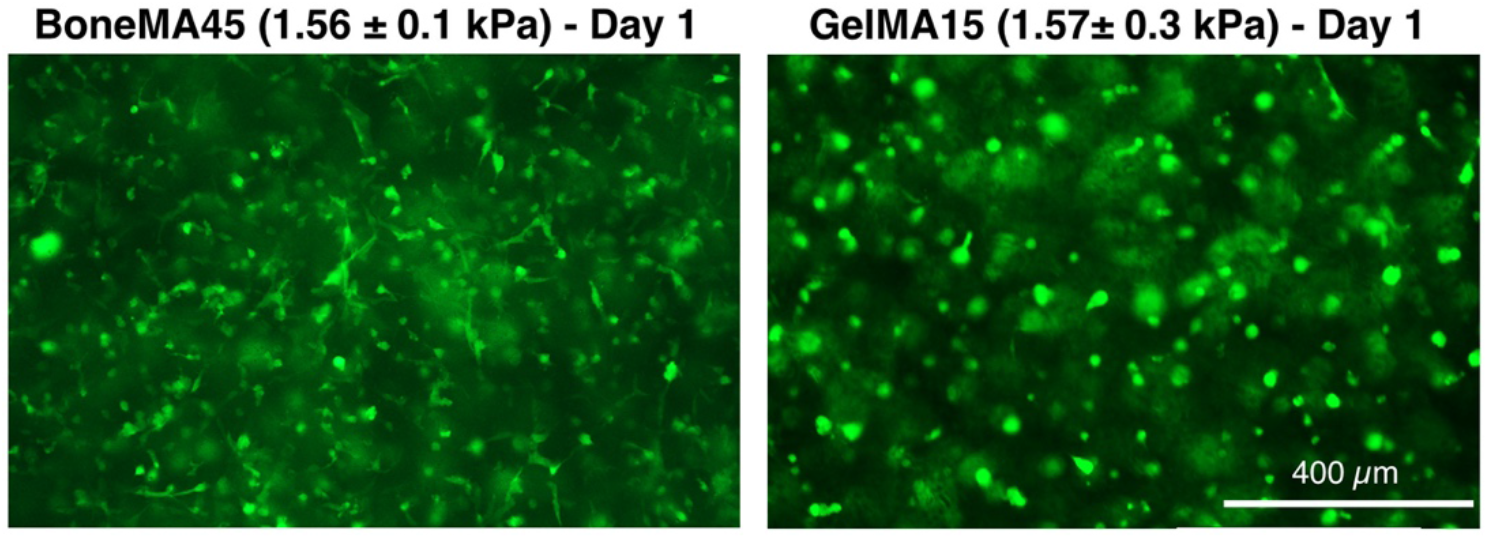
BoneMA hydrogels (with an elastic modulus of 1.56 ±0.1 kPa) were more vasculogenic than GelMA hydrogels of similar stiffness (elastic modulus 1.57 ±0.3 kPa) with visibly better vascular network formation by HUVECs in BoneMA within one day of encapsulation. (Scale bar – 400 μm)

Quantification of vascular network formation (Figure 6) showed that the number of vascular segments was, in general, higher in BoneMA samples compared to that in GelMA. In particular, there was a significant difference between BoneMA exposed to 45 seconds of photopolymerization, and GelMA exposed for 30 seconds, as the number of segments increased in BoneMA by a factor of 1.6 (*p* < 0.001).

**Figure 6.**
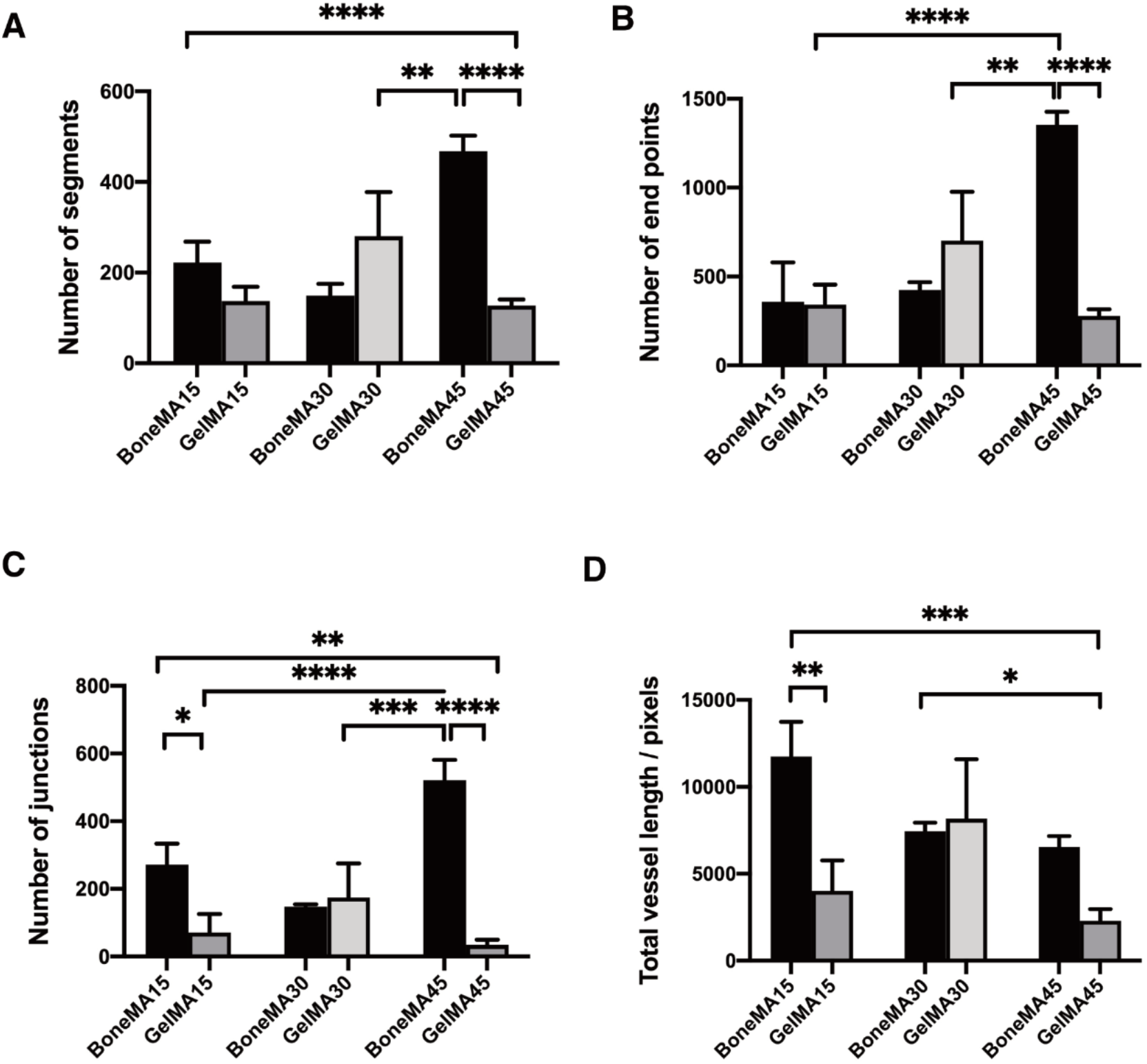
Characterization of vascular networks in BoneMA and GelMA after 3 days. Quantitative analysis of the vessel networks measured the (A) segments, (B) end points, (C) junctions, and (D) vessel length. The total vessel length after 3 days was significantly higher in BoneMA15 than GelMA15 (*p* < 0.01), while there was no statistically significant difference among BoneMA30 and GelMA30, and BoneMA45 and GelMA45. (\**p* < 0.05; \*\**p* < 0.01; \*\*\**p* < 0.001; \*\*\*\**p* < 0.0001).

Similarly, there was a significant difference in the number of segments between samples BoneMA15 versus GelMA45 (*p* < 0.0001), and BoneMA45 versus GelMA45 (*p* < 0.0001). The number of endpoints, which represents the number of sprouts found at the ends of the vessel as well as the tip in side-sprouting networks, was also higher in BoneMA, suggesting an active network formation in the hydrogel. We also found a significant difference among GelMA30 versus BoneMA45 (*p* < 0.01), GelMA15 versus BoneMA45 (*p*< 0.0001), and BoneMA45 versus GelMA45 (*p* < 0.0001), where the number of junctions was higher in BoneMA than GelMA. The total vessel length was also generally higher in BoneMA hydrogels. For instance, the total vessel length for BoneMA15 increased by a factor of 5.1 (*p* < 0.001) in comparison to GelMA45, and 2.9 (*p* < 0.005) for GelMA15. All these quantitative data show a very active network forming process occurring in BoneMA when compared to GelMA.

Lastly, fluorescence microscopy images of bioprinted patterns using BoneMA encapsulating HMSCs and immunostained for F-Actin (Figure 7) show a variety of geometrical patterns of different sizes. Micropatterns that ranged from 600 μm (square) to 2.5 mm (OHSU logo) could be bioprinted consistently in as little as 15 sec. Of note, these microgels can be bioprinted with the exact sample printing parameters optimized for vasculature formation, as shown in previous images, thereby allowing for fabrication of pre-vascularized microscaffolds at a time. We also printed the positive and negative features of BoneMA without cells with dimensions ranging from 500 μm to 1000 μm (Supplementary Figure S4). Importantly, these microgels can be injected through a standard syringe needle, thus forming a straightforward platform for the fabrication of pre-vascularized injectable microgels with advantageous vasculogenic properties.

**Figure 7.**
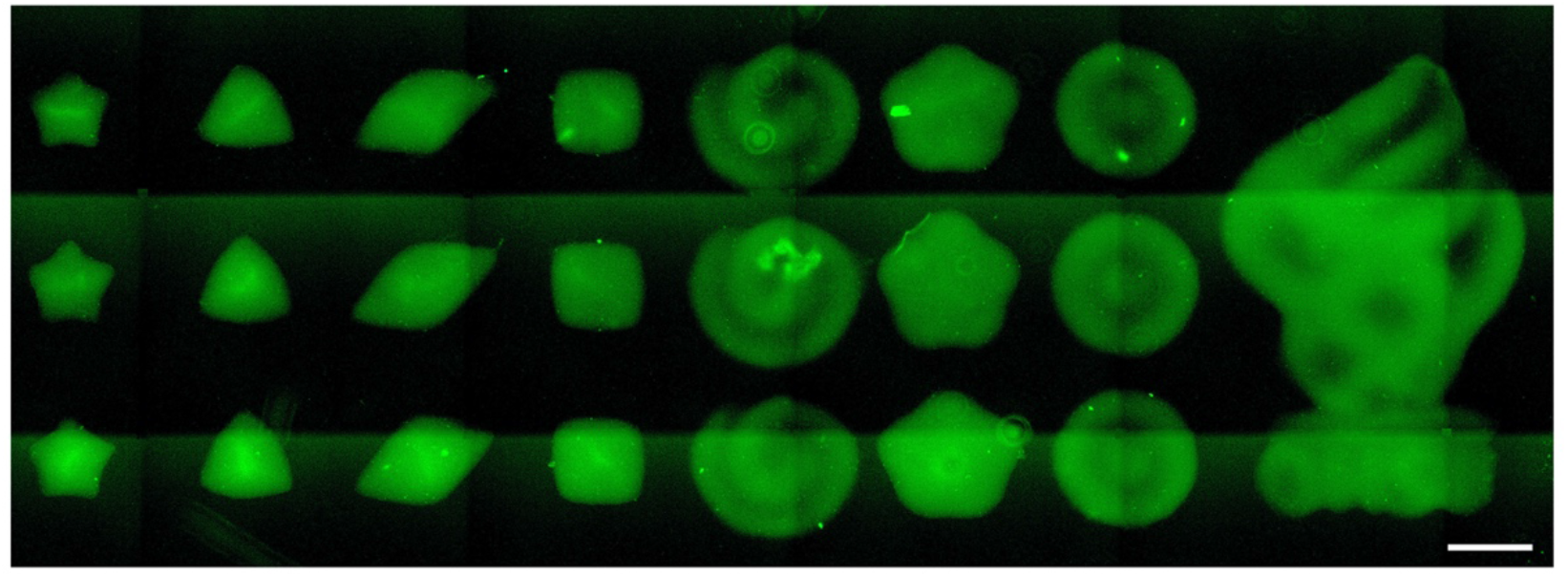
Representative fluorescence microscopy image demonstrating the range of patterns and geometries achieved by printing HMSC-laden BoneMA with a DLP printer (Ember). The cells were stained with F-Actin (green) after printing to enable confirmation of the shape and structure of the printed constructs through fluorescence microscopy. (Scale – 750 μm)

## 4. Discussion

In this paper, we demonstrate the synthesis and characterization of a photocrosslinkable ECM-derived hydrogel utilizing the native human bone matrix as a starting point, which we refer to as methacrylated bone, or BoneMA. BoneMA hydrogels showed superior vasculogenic potential when compared to GelMA and also enhanced the ability to be bioprinted into microscale geometries, both with and without cells.

### 4.1. Synthesis of BoneMA

Our first goal was to synthesize a new hydrogel biomaterial from a decellularized and demineralized bone matrix with controllable physical properties. As native collagen is the principal component of the bone ECM, demineralized bone matrix materials share the drawback of a lack of control over mechanical properties. From an engineering point of view, this is a constraint in regulating tissue mechanics [23], which is an essential factor in determining cell behavior and differentiation. The purpose of protein methacrylation is to endow the matrix with photocrosslinkable moieties that allow for the control of the final mechanical properties of the methacrylated biomaterial [24]. Methacrylated gelatin (GelMA) was used as a control to compare some basic physical and biological properties of BoneMA relative to a well-established matrix material. In the GelMA synthesis, pH is maintained at 7.4 by the phosphate buffer, which introduces methacryloyl groups to the reactive amine and hydroxyl groups of the amino acid residues [25] (Fig. 1B). The main limitation in the above process is that the referred pH limits the reactivity of amine and hydroxyl groups. To tackle this problem, we modified an improved synthesis protocol [26], where a higher pH (9.0) is maintained during the reaction using carbonate bi-carbonate as a buffer to keep the isoelectric point (IEP) of the bone proteins high, keeping the free amino groups of lysine neutral and allowing for it to react with methacrylic anhydride.

When comparing BoneMA against GelMA, it is important to remember that BoneMA is a multi-component hydrogel that carries the proteolytic by-products of pepsinized collagen and resulting peptides of a wide range of non-collagenous proteins. Single component hydrogels, such as GelMA, methacrylated hyaluronic acid (meHA) or methacrylated tropoelastin (MeTro) have only one major component like gelatin, HA, and tropoelastin, respectively [27]. The methacrylation of these single component hydrogels also uses a similar synthesis route. For example, the methacrylation of tropoelastin was done on the lysine residues in the structure, whereas in meHA, the carboxyl and primary hydroxyl groups of HA underwent functionalization [18, 28]. Therefore, the differences in physical properties that are identified in GelMA and BoneMA may be attributed to the heterogeneity in the molecular network that is formed in the cross-linked bone peptides (of different lengths and types) versus the more homogenous and controlled one-component matrix of GelMA. Finally, it is worth noting that like other methacrylated matrix hydrogels, the mechanical properties of BoneMA can also be tailored by changing the methacrylate concentration, photoinitiator concentration, exposure time and/or combination of the parameters, which is advantageous in controlling the stem cell microenvironment and allowing for in-situ photopolymerization using light sources that are used clinically, as we reported previously [29]. Moreover, as in previous reports, we show that BoneMA can be photocured using LAP as the photoinitiator, which allows it to be crosslinked in the visible wavelength region (400-420 nm) [30] (Fig. 1B).

The rheological studies revealed relevant information regarding the photocrosslinking process of BoneMA (Supplementary Figure S5A). The gradual slope of the curve indicates that the modulus increased with an increase in photocrosslinking time, as expected. GelMA too showed a gradual slope in shear modulus, but at a higher rate than BoneMA (Supplementary Figure S5B). This indicates that BoneMA is a much softer material than GelMA, which makes for a desirable material when lower stiffness is needed, such as in vascular regeneration [29]. With respect to bioprinting, specifically, the storage modulus of the material also plays an important role. It is reported that when a hydrogel has a storage modulus in the order of 100-1000 Pa, it is also suitable for extrusion bioprinting [31], although the details of such a process fall beyond the scope of the current paper.

### 4.2. BoneMA enhances endothelial cell vasculogenesis

Next, our goal was to investigate how BoneMA affects endothelial cells during the process of vascular morphogenesis, which is an essential process for tissue regeneration [32]. To test this, HUVECs were encapsulated in BoneMA and GelMA hydrogels (**Figure 4**). GelMA is an established biomaterial in promoting vascular network formation using endothelial cells [33], therefore it was chosen as a control. Surprisingly, we found that BoneMA formed well-established vascular networks in 48 hours in all samples, irrespective of crosslinking time. During the same period, in GelMA, HUVECs only started to extend their filopodia for establishing cell-cell contact. This faster network formation established the inherent ability of BoneMA to induce vascular morphogenesis. The network formation in BoneMA could be seen on day 2, while GelMA took three days to form similar networks. Based on the crosslinking density of the hydrogel, the steps involved in network formation, such as matrix degradation, cell migration, and cell-cell contact appear to be somewhat affected.

Since the stiffness of cell-laden hydrogels has previously been linked to the ability of HUVECs to form microvascular networks in GelMA [29], where hydrogels of elastic modulus below 2.5 kPa showed enhanced vasculature formation, we hypothesized that the intrinsically lower mechanical properties of BoneMA could be the reason for the improved results. To test this hypothesis, we compared BoneMA and GelMA of similar elastic moduli cultured with HUVECS in similar conditions. Interestingly, GelMA and BoneMA, which had an equivalent elastic modulus of 1.56 ±0.1 kPa, showed visibly faster network formation in BoneMA (**Figure 5**). This suggests that despite the mechanical properties, BoneMA has an inherent ability to promote faster vascular network formation in-vitro. A rational explanation for such an improved vasculature formation in-vitro can be attributed to the host of pro-angiogenic peptides that have been reported to be present in the bone ECM, and are likely to survive the synthesis process, although we have not quantified these molecules [34].

### 4.3. BoneMA as a bioink

Lastly, in order to further demonstrate the versatility of BoneMA as a bioink for DLP bioprinting of microscale geometries [35], we printed structures leading to the formation of injectable ‘granular’ microgels. Microgels are a special class of materials that have gained attention in fabricating complex tissues. Microgels are crosslinked polymer networks in the micron range, which has considerable advantages over bulk hydrogels in the area of controlled release of drugs and protein, hydrolytic degradation, personalized medicine screening and microscale tissue engineering [36, 37]. In microscale tissue engineering, microgels may be used as building blocks for the bottom-up building of hetero-architecture to mimic complex tissues. Shape is a crucial factor in microgel geometry to create centimeter-scale tissue-like structures. For example, tightly packed cell-laden tissue-like structures were created through a bottom-up self-assembly in a multiphase environment (liquid-air system) [38]. Hexagonal cell-laden microgels formed the tissue-like structures through an interface-directed assembly process, while more complex building blocks like lock-and-key microgels allowed greater control over self-assembly. Since cell-laden BoneMA allows printing in multiple complex shapes (Figure 7), similar lock-and-key shaped cell-laden microgels can be printed to create complex shapes and tissues. Also, culturing BoneMA microgels with two or more cells can further address the complexity of building tissues. Therefore, we foresee BoneMA microgels as an effective ECM-based building block for bottom-up manufacturing of complex tissues. Injectability of microgels is another parameter that plays a major role in therapeutic delivery for treating site-specific tissue damage like myocardial infarction [39], enhancing neovascularization [40] and cellular differentiation [41]. The injectability of BoneMA microgels opens up venues for investigations in basic biology and clinically-oriented minimally invasive implantation procedures (Supplementary Figure S6).

## Conclusions

In summary, our data suggest the successful preparation of a photocrosslinkable hydrogel from demineralized and decellularized human bone. One of the key findings is that, not only BoneMA supports vascularization of endothelial cells, but it also initiates vascularization more quickly than common hydrogels of comparable chemistry, such as GelMA. Further, we envision that since BoneMA has an excellent print fidelity in the micro dimensions, it may facilitate the production of vascularized microgels in high-throughput. We have also established BoneMA as a bioink, and this will set a new paradigm in the development of photocrosslinkable human ECM-derived bioinks in the field of bioprinting.

## Acknowledgments

This project was supported by funding from the National Institute of Dental and Craniofacial Research (R01DE026170 and 3R01DE026170-03S1 to LEB), Oregon Clinical & Translational Research Institute (OCTRI) - Biomedical Innovation Program (BIP), IADR-GSK Innovation in Oral Care Awards, Michigan-Pittsburgh-Wyss Resource Center - Regenerative Medicine Resource Center (MPW-RM), and the OHSU Fellowship for Diversity and Inclusion in Research (OHSU-OFDIR to CMF).

---

**Supplementary Figure S1.** Physical characterization of GelMA. (A) The elastic modulus of GelMA hydrogels increased as a function of crosslinking duration starting from 1.5 kPa at 15 sec to 3 kPa and 4.3 kPa at 30 and 45 sec, respectively. Meanwhile, apparent pore size as evidenced by SEM images of (B) freeze dried and (C) critical point dried GelMA samples showed a decrease in pore size from (i) GelMA hydrogels crosslinked for 15 sec to those that were crosslinked for (ii) 30 and (iii) 45 sec. (Scale – 5 μm). (\**p* < 0.05; \*\**p* < 0.005; \*\*\**p* < 0.001)

**Supplementary Figure S2**. A) Apparent diffusion of rhodamine dye, as measured by its fluorescence, was higher in BoneMA hydrogels polymerized for 15 sec in comparison with those that were polymerized for 30 and 45 sec, suggesting that the less crosslinked hydrogels were more permeable. (B) For the swelling analysis, BoneMA hydrogel discs (n = 6) were stored in DPBS at 37 C for a day, blot dried, and the wet swollen weight was recorded. After weighing, the samples were lyophilized, and the dried samples were weighed again to record their corresponding dry weight. The equilibrium swelling ratio was calculated as the ratio of wet mass to the dry mass of the hydrogel. The swelling properties of all the samples were identical over the given time period. Scale bar – 50 μm.

**Supplementary Figure S3.** Cytocompatibility of GelMA hydrogels. Representative images of hDPSCs encapsulated in GelMA hydrogels crosslinked for 15, 30, and 45 secs and stained for live (blue) and dead (green) cells at 1, 3 and 7 day time points showed a high degree of viability across all conditions and time points. Scale bar - 400 μm.

**Supplementary Figure S4**. Both positive and negative features ranging from 500 - 1000 μm in width were printed using BoneMA after a crosslinking time of 45 seconds. The resolution for the negative was better than positive for the 500 μm square print.

**Supplementary Figure S5**. Shear storage modulus of BoneMA and GelMA. A) The shear storage modulus of BoneMA increased during the first 100 seconds of crosslinking. During this period, the shear storage modulus reached approximately 500 Pa, and beyond 100 seconds, the value reached approximately 750 Pa, where the curve remained a constant after 150 seconds. B) GelMA had a sharp increase in storage modulus during the first 50 seconds, and the value reached approximately 2 kPa. The curve hit a constant at 125 seconds when the shear storage modulus reached a value of approximately 2.5 kPa.

**Supplementary Figure S6**. Injectability of BoneMA microgels. (**A-F**) Image sequence of BoneMA microgel delivery through a gauge 18 needle. The microgels were easily injected through the syringe and remains confined within the mold.

## References

[1] C. Frantz, K.M. Stewart, V.M. Weaver, The extracellular matrix at a glance, Journal of Cell Science 123(24) (2010) 4195–4200

[2] M. Chiquet, Regulation of extracellular matrix gene expression by mechanical stress, Matrix Biology 18(5) (1999) 417–426.

[3] D.A. Siwik, P.J. Pagano, W.S. Colucci, Oxidative stress regulates collagen synthesis and matrix metalloproteinase activity in cardiac fibroblasts, American Journal of Physiology-Cell Physiology 280(1) (2001) C53–C60.

[4] T. Barber, G. Esteban-Pretel, M. Marín, J. Timoneda, Vitamin A Deficiency and Alterations in the Extracellular Matrix, Nutrients 6(11) (2014) 4984–5017.

[5] W.P. Daley, S.B. Peters, M. Larsen, Extracellular matrix dynamics in development and regenerative medicine, Journal of Cell Science 121(3) (2008) 255–264.

[6] J. Sottile, Regulation of angiogenesis by extracellular matrix, Biochimica et Biophysica Acta (BBA) - Reviews on Cancer 1654(1) (2004) 13–22.

[7] F. Gattazzo, A. Urciuolo, P. Bonaldo, Extracellular matrix: A dynamic microenvironment for stem cell niche, Biochimica et Biophysica Acta (BBA) - General Subjects 1840(8) (2014) 2506–2519.

[8] H. Järveläinen, A. Sainio, M. Koulu, T.N. Wight, R. Penttinen, Extracellular Matrix Molecules: Potential Targets in Pharmacotherapy, Pharmacol. Rev. 61(2) (2009) 198–223.

[9] E.L. Smith, J.M. Kanczler, D. Gothard, C.A. Roberts, J.A. Wells, L.J. White, O. Qutachi, M.J. Sawkins, H. Peto, H. Rashidi, L. Rojo, M.M. Stevens, A.J. El Haj, F.R.A.J. Rose, K.M. Shakesheff, R.O.C. Oreffo, Evaluation of skeletal tissue repair, Part 1: Assessment of novel growth-factor-releasing hydrogels in an ex vivo chick femur defect model, Acta Biomater. 10(10) (2014) 4186–4196.

[10] E.L. Smith, J.M. Kanczler, D. Gothard, C.A. Roberts, J.A. Wells, L.J. White, O. Qutachi, M.J. Sawkins, H. Peto, H. Rashidi, L. Rojo, M.M. Stevens, A.J. El Haj, F.R.A.J. Rose, K.M. Shakesheff, R.O.C. Oreffo, Evaluation of skeletal tissue repair, Part 2: Enhancement of skeletal tissue repair through dual-growth-factor-releasing hydrogels within an ex vivo chick femur defect model, Acta Biomater. 10(10) (2014) 4197–4205.

[11] J. Wu, Q. Ding, A. Dutta, Y. Wang, Y.-h. Huang, H. Weng, L. Tang, Y. Hong, An injectable extracellular matrix derived hydrogel for meniscus repair and regeneration, Acta Biomater. 16 (2015) 49–59.

[12] H. Ghuman, A.R. Massensini, J. Donnelly, S.-M. Kim, C.J. Medberry, S.F. Badylak, M. Modo, ECM hydrogel for the treatment of stroke: Characterization of the host cell infiltrate, Biomaterials 91 (2016) 166–181.

[13] D. Tukmachev, S. Forostyak, Z. Koci, K. Zaviskova, I. Vackova, K. Vyborny, I. Sandvig, A. Sandvig, C.J. Medberry, S.F. Badylak, E. Sykova, S. Kubinova, Injectable Extracellular Matrix Hydrogels as Scaffolds for Spinal Cord Injury Repair, Tissue Engineering Part A 22(3-4) (2016) 306–317.

[14] D. Chaimov, L. Baruch, S. Krishtul, I. Meivar-levy, S. Ferber, M. Machluf, Innovative encapsulation platform based on pancreatic extracellular matrix achieve substantial insulin delivery, Journal of Controlled Release 257 (2017) 91–101.

[15] M.T. Spang, K.L. Christman, Extracellular matrix hydrogel therapies: In vivo applications and development, Acta Biomater. 68 (2018) 1–14.

[16] J.W. Nichol, S.T. Koshy, H. Bae, C.M. Hwang, S. Yamanlar, A. Khademhosseini, Cell-laden microengineered gelatin methacrylate hydrogels, Biomaterials 31(21) (2010) 5536–44.

[17] K. Yue, G. Trujillo-de Santiago, M.M. Alvarez, A. Tamayol, N. Annabi, A. Khademhosseini, Synthesis, properties, and biomedical applications of gelatin methacryloyl (GelMA) hydrogels, Biomaterials 73 (2015) 254–271.

[18] J.A. Burdick, C. Chung, X. Jia, M.A. Randolph, R. Langer, Controlled Degradation and Mechanical Behavior of Photopolymerized Hyaluronic Acid Networks, Biomacromolecules 6(1) (2005) 386–391.

[19] M.J. Sawkins, W. Bowen, P. Dhadda, H. Markides, L.E. Sidney, A.J. Taylor, F.R.A.J. Rose, S.F. Badylak, K.M. Shakesheff, L.J. White, Hydrogels derived from demineralized and decellularized bone extracellular matrix, Acta Biomater. 9(8) (2013) 7865–7873.

[20] N. Monteiro, G. Thrivikraman, A. Athirasala, A. Tahayeri, C.M. França, J.L. Ferracane, L.E. Bertassoni, Photopolymerization of cell-laden gelatin methacryloyl hydrogels using a dental curing light for regenerative dentistry, Dent. Mater. 34(3) (2018) 389–399.

[21] V.A. Simms, R. Bicknell, V.L. Heath, Development of an ImageJ-based method for analysing the developing zebrafish vasculature, Vascular Cell 9(1) (2017) 2.

[22] A. Lesman, D. Rosenfeld, S. Landau, S. Levenberg, Mechanical regulation of vascular network formation in engineered matrices, Advanced Drug Delivery Reviews 96 (2016) 176–182.

[23] I.D. Gaudet, D.I. Shreiber, Characterization of Methacrylated Type-I Collagen as a Dynamic, Photoactive Hydrogel, Biointerphases 7(1) (2012).

[24] E. Hachet, H. Van Den Berghe, E. Bayma, M.R. Block, R. Auzély-Velty, Design of Biomimetic Cell-Interactive Substrates Using Hyaluronic Acid Hydrogels with Tunable Mechanical Properties, Biomacromolecules 13(6) (2012) 1818–1827.

[25] A.I. Van Den Bulcke, B. Bogdanov, N. De Rooze, E.H. Schacht, M. Cornelissen, H. Berghmans, Structural and Rheological Properties of Methacrylamide Modified Gelatin Hydrogels, Biomacromolecules 1(1) (2000) 31–38.

[26] H. Shirahama, B.H. Lee, L.P. Tan, N.-J. Cho, Precise Tuning of Facile One-Pot Gelatin Methacryloyl (GelMA) Synthesis, Scientific Reports 6(1) (2016).

[27] N. Annabi, S.M. Mithieux, P. Zorlutuna, G. Camci-Unal, A.S. Weiss, A. Khademhosseini, Engineered cell-laden human protein-based elastomer, Biomaterials 34(22) (2013) 5496–5505.

[28] M. Yamamoto, M.T. Poldervaart, B. Goversen, M. de Ruijter, A. Abbadessa, F.P.W. Melchels, F.C. Öner, W.J.A. Dhert, T. Vermonden, J. Alblas, 3D bioprinting of methacrylated hyaluronic acid (MeHA) hydrogel with intrinsic osteogenicity, PLoS One 12(6) (2017).

[29] N. Monteiro, W. He, C.M. Franca, A. Athirasala, L.E. Bertassoni, Engineering Microvascular Networks in LED Light-Cured Cell-Laden Hydrogels, ACS Biomaterials Science & Engineering 4(7) (2018) 2563–2570.

[30] B.D. Fairbanks, M.P. Schwartz, C.N. Bowman, K.S. Anseth, Photoinitiated polymerization of PEG-diacrylate with lithium phenyl-2,4,6-trimethylbenzoylphosphinate: polymerization rate and cytocompatibility, Biomaterials 30(35) (2009) 6702–6707.

[31] N. Ashammakhi, S. Ahadian, C. Xu, H. Montazerian, H. Ko, R. Nasiri, N. Barros, A. Khademhosseini, Bioinks and bioprinting technologies to make heterogeneous and biomimetic tissue constructs, Materials Today Bio 1 (2019) 100008.

[32] X. Chen, A.S. Aledia, S.A. Popson, L. Him, C.C.W. Hughes, S.C. George, Rapid Anastomosis of Endothelial Progenitor Cell–Derived Vessels with Host Vasculature Is Promoted by a High Density of Cotransplanted Fibroblasts, Tissue Engineering Part A 16(2) (2010) 585–594.

[33] Y.-C. Chen, R.-Z. Lin, H. Qi, Y. Yang, H. Bae, J.M. Melero-Martin, A. Khademhosseini, Functional Human Vascular Network Generated in Photocrosslinkable Gelatin Methacrylate Hydrogels, Adv. Funct. Mater. 22(10) (2012) 2027–2039.

[34] A. Neve, F.P. Cantatore, N. Maruotti, A. Corrado, D. Ribatti, Extracellular Matrix Modulates Angiogenesis in Physiological and Pathological Conditions, BioMed Research International 2014 (2014) 1–10.

[35] D. Choudhury, H.W. Tun, T. Wang, M.W. Naing, Organ-Derived Decellularized Extracellular Matrix: A Game Changer for Bioink Manufacturing?, Trends in Biotechnology 36(8) (2018) 787–805.

[36] J.K. Oh, R. Drumright, D.J. Siegwart, K. Matyjaszewski, The development of microgels/nanogels for drug delivery applications, Prog. Polym. Sci. 33(4) (2008) 448–477.

[37] W. Yang, H. Yu, G. Li, Y. Wang, L. Liu, High-Throughput Fabrication and Modular Assembly of 3D Heterogeneous Microscale Tissues, Small 13(5) (2017).

[38] B. Zamanian, M. Masaeli, J.W. Nichol, M. Khabiry, M.J. Hancock, H. Bae, A. Khademhosseini, Interface-Directed Self-Assembly of Cell-Laden Microgels, Small 6(8) (2010) 937–944.

[39] M.H. Chen, J.J. Chung, J.E. Mealy, S. Zaman, E.C. Li, M.F. Arisi, P. Atluri, J.A. Burdick, Injectable Supramolecular Hydrogel/Microgel Composites for Therapeutic Delivery, Macromolecular Bioscience 19(1) (2019).

[40] P.-H. Kim, H.-G. Yim, Y.-J. Choi, B.-J. Kang, J. Kim, S.-M. Kwon, B.-S. Kim, N.S. Hwang, J.-Y. Cho, Injectable multifunctional microgel encapsulating outgrowth endothelial cells and growth factors for enhanced neovascularization, Journal of Controlled Release 187 (2014) 1–13.

[41] Y. Hou, W. Xie, K. Achazi, J.L. Cuellar-Camacho, M.F. Melzig, W. Chen, R. Haag, Injectable degradable PVA microgels prepared by microfluidic technology for controlled osteogenic differentiation of mesenchymal stem cells, Acta Biomater. 77 (2018) 28–37.

