## Supplementary figures for "BoneMA – Synthesis and Characterization of a Methacrylated Bone-derived Hydrogel for Bioprinting of Vascularized Tissues"

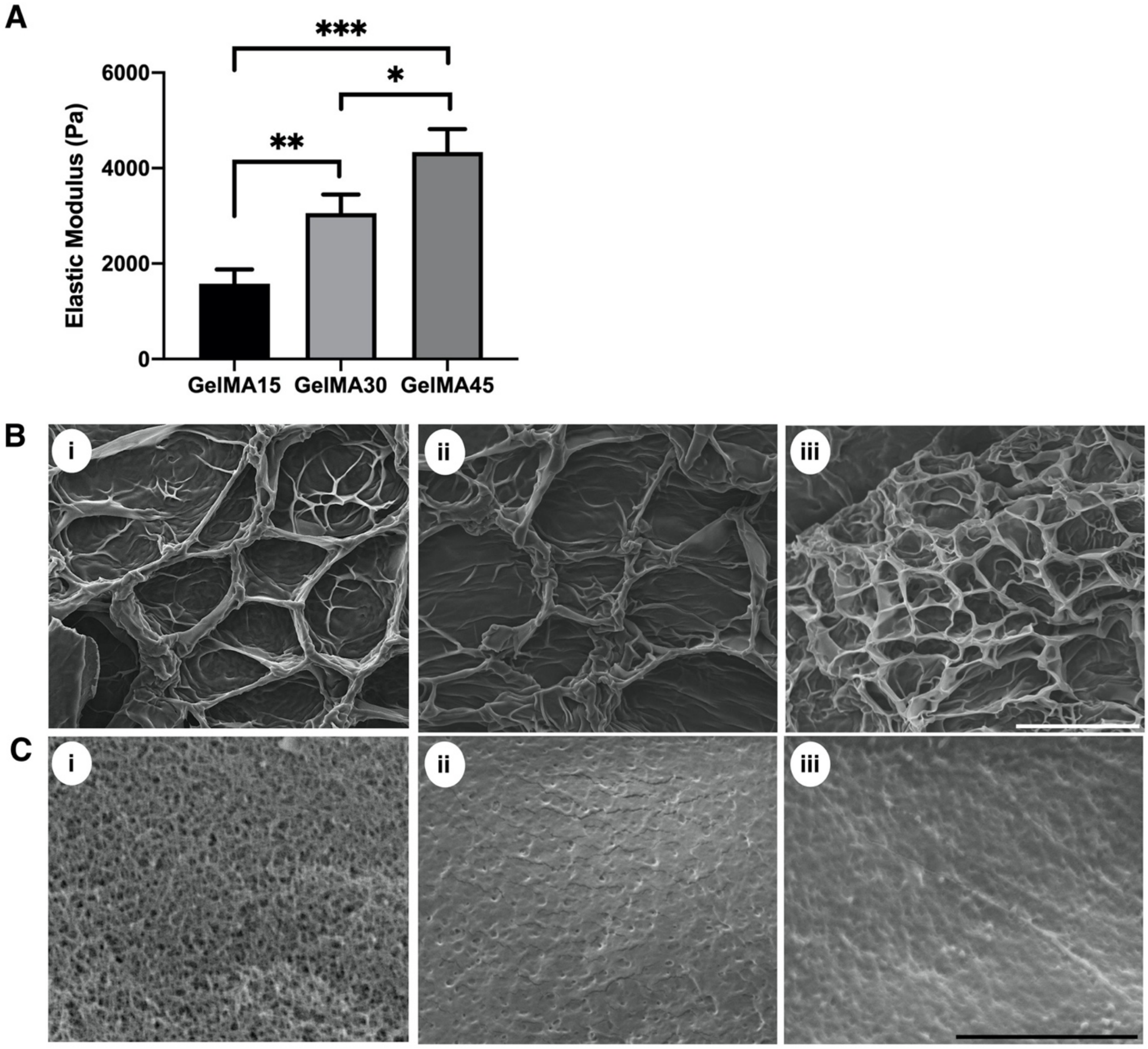


**Supplementary figure S1.** Physical characterization of GelMA. (A) The elastic modulus of GelMA hydrogels increased as a function of crosslinking duration starting from 1.5 kPa at 15 sec to 3 kPa and 4.3 kPa at 30 and 45 sec, respectively. Meanwhile, apparent pore size as evidenced by SEM images of (B) freeze dried and (C) critical point dried GelMA samples showed a decrease in pore size from (i) GelMA hydrogels crosslinked for 15 sec to those that were crosslinked for (ii) 30 and (iii) 45 sec. (Scale – 5 µm) (*$p$ < 0.05; **$p$ < 0.005; ***$p$ < 0.001).

A

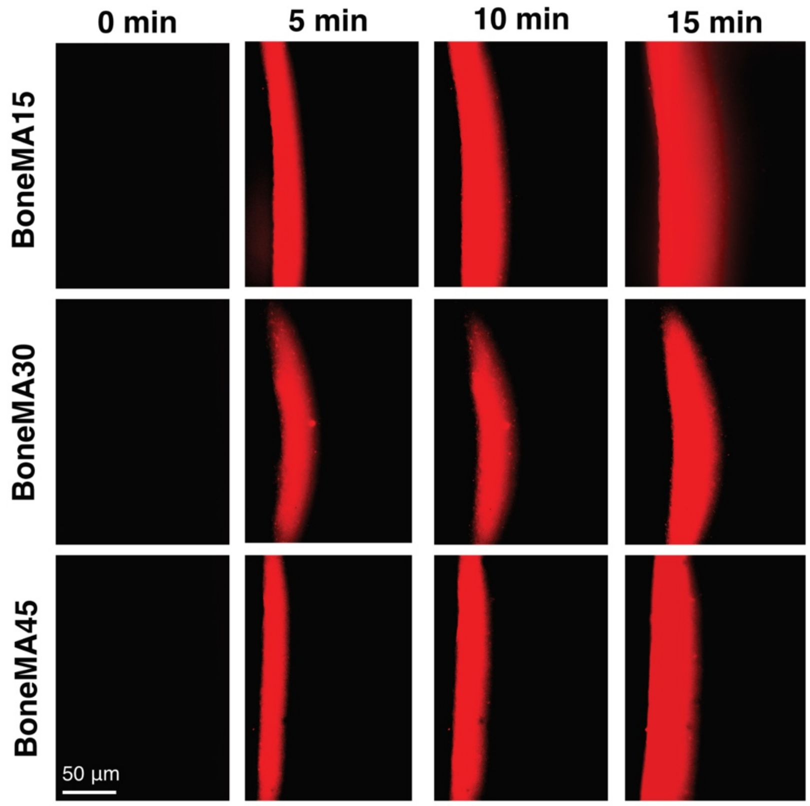


B

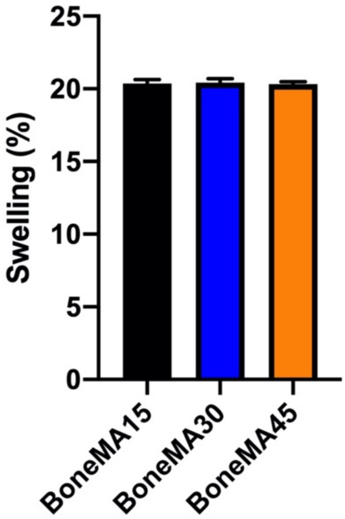


**Supplementary figure S2.** (A) Apparent diffusion of rhodamine dye, as measured by its fluorescence, was higher in BoneMA hydrogels polymerized for 15 sec in comparison with those that were polymerized for 30 and 45 sec, suggesting that the less crosslinked hydrogels were more permeable. (B) For the swelling analysis, BoneMA hydrogel discs (n = 6) were stored in DPBS at 37 °C for a day, blot dried, and the wet swollen weight was recorded. After weighing, the samples were lyophilized, and the dried samples were weighed again to record their corresponding dry weight. The equilibrium swelling ratio was calculated as the ratio of wet mass to the dry mass of the hydrogel. The swelling properties of all the samples were identical over the given time period. Scale bar – 50 µm

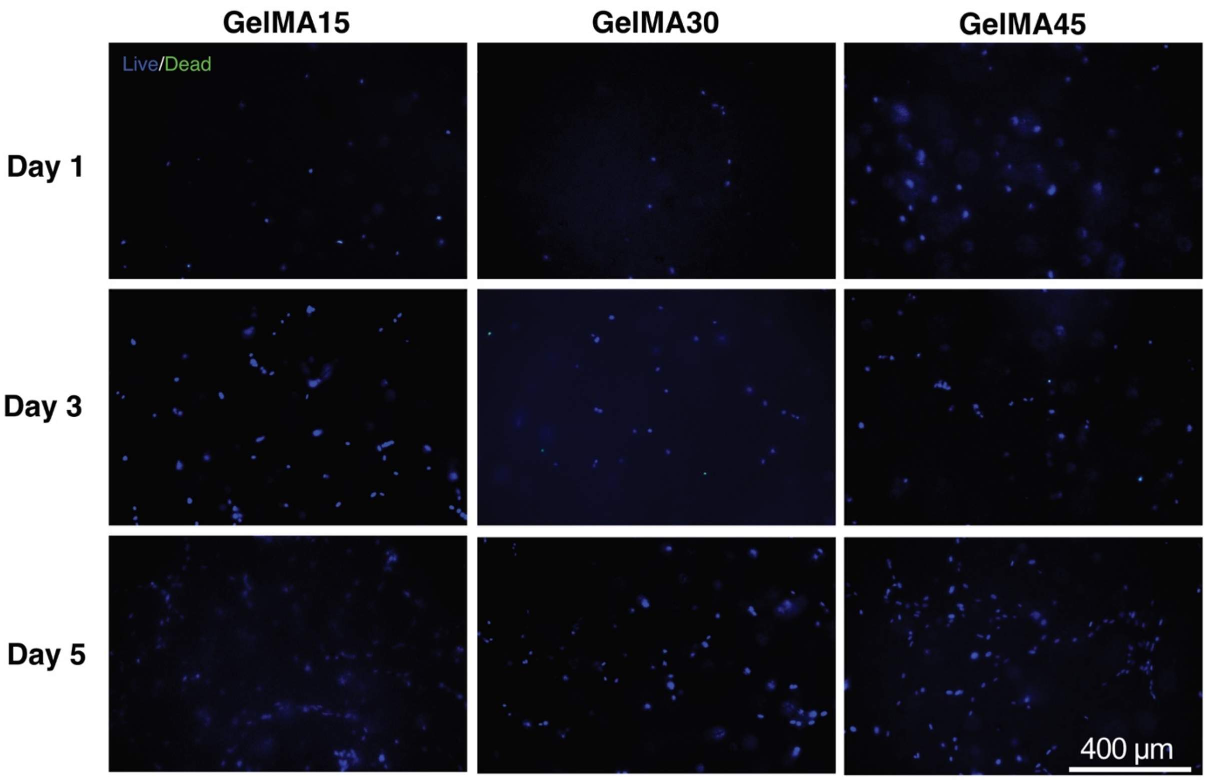


**Supplementary figure S3**. Cytocompatibility of GelMA hydrogels. Representative images of hDPSCs encapsulated in GelMA hydrogels crosslinked for 15, 30, and 45 secs and stained for live (blue) and dead (green) cells at 1, 3 and 7 day time points showed a high degree of viability across all conditions and time points. Scale bar - 400 µm

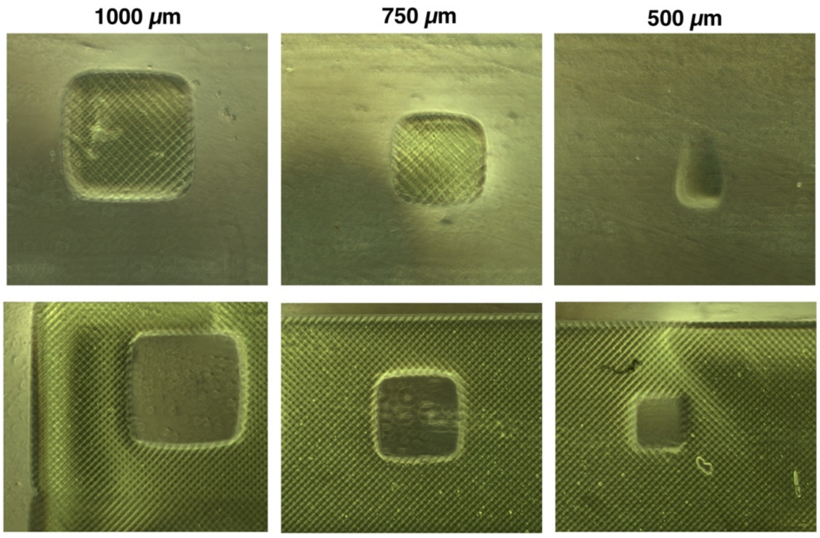


**Supplementary figure S4.** Both positive and negative features ranging from 500 - 1000 µm in width were printed using BoneMA after a crosslinking time of 45 seconds. The resolution for the negative was better than positive for the 500 µm square print.

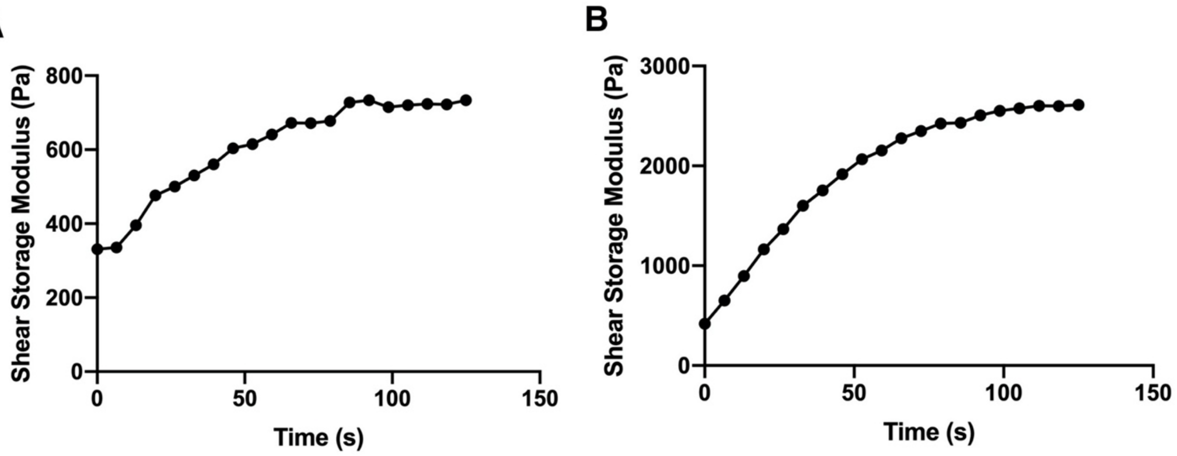


**Supplementary figure S5**. Shear storage modulus of BoneMA and GelMA. **A**) The shear storage modulus of BoneMA increased during the first 100 seconds of crosslinking. During this period, the shear storage modulus reached approximately 500 Pa, and beyond 100 seconds, the value reached approximately 750 Pa, where the curve remained a constant after 150 seconds. **B**) GelMA had a sharp increase in storage modulus during the first 50 seconds, and the value reached approximately 2 kPa. The curve hit a constant at 125 seconds when the shear storage modulus reached a value of approximately 2.5 kPa.

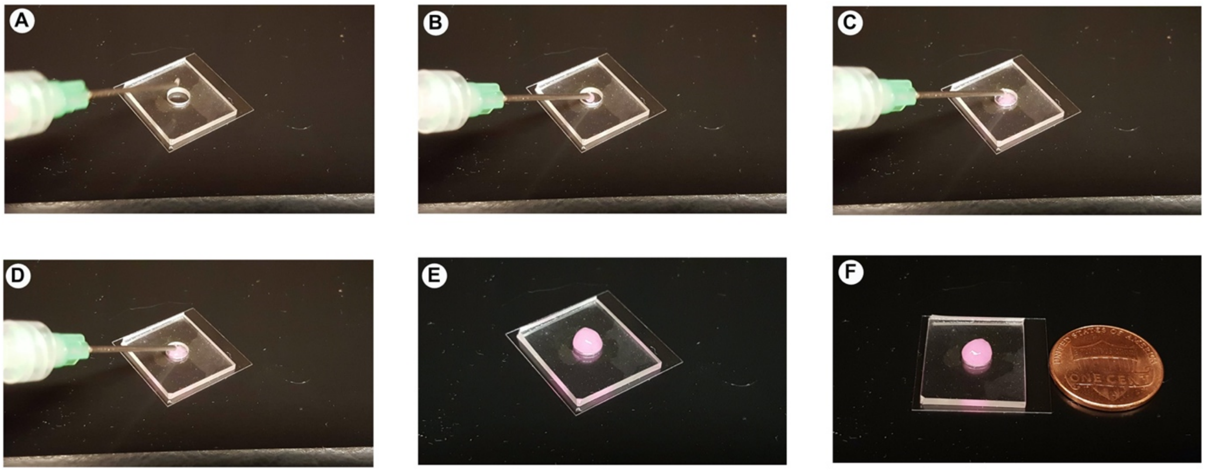


**Supplementary figure S6**. Injectability of BoneMA microgels. (**A-F**) Image sequence of BoneMA microgel delivery through a gauge 18 needle. The microgels were easily injected through the syringe and remains confined within the mold.

### Supplementary methods

*S1. Preparation of gelatin methacryloyl (GelMA)*

Gelatin methacryloyl (GelMA) was used as a control to compare against the biological properties of BoneMA. GelMA was synthesized as per the protocol described by Nichol *et al.* [16]. Porcine skin type A gelatin (10% w/v) (Sigma, St Louis, MO, USA) was dissolved in Dulbecco's phosphate buffered saline (DPBS, Sigma) warmed to 50 °C to which, 8% (v/v) methacrylic anhydride (Sigma) was added dropwise and allowed to react for 2 h. Next, the solution was diluted 5x times with DPBS and dialyzed with 12-14 kDa dialysis tubing against warm distilled water (45 ± 5 °C) for 5 days. The warm water was changed two times a day for 5 days. The resulting methacrylated prepolymer was lyophilized and stored at room temperature until further use. For GelMA sample synthesis, GelMA was crosslinked for 15 sec, 30 sec, and 45 sec using the bioprinter as described previously, and the resultant constructs are identified as GelMA15, GelMA30, and GelMA45 respectively. The GelMA samples were treated similarly to BoneMA for SEM and live/dead analysis.
